# Identification of target candidate in Polycystic ovarian syndrome and invitro evaluation of therapeutic activity of the designed RNA Aptamer

**DOI:** 10.1101/603357

**Authors:** Manibalan Subramanian, Shobana Ayyachamy, Kiruthika Manickam, Swathi Madasamy, Venkatalakshmi Renganathan, Anant Achary, Thirukumaran Kandasamy, K Suhasini, Sharon Roopathy

## Abstract

Prevalence of poly cystic ovary syndrome has been gradually increasing among adult females and Laparoscopy drilling on the ovary is only available temporary solution with high incidence of reoccurrence. Confidential gene disease association studies combined with various graph theory analysis on the biological protein interaction network of this syndrome has resulted, the 15 genes are overexpressed as central nodes among 434 proteins of disease specific proteome network. Through the Intensive Structural and functional prioritization analysis we have identified S100A8, calprotectin is the ideal drug target protein. In this research, we have constructed RNA library of consensus DNA sequence of Glucocorticoid Response Element (GRE) and screened the best RNA Aptamer fragment which competitively binds with minimal energy to inhibit the cell migration activity of S100A8. In order to prove this computational research, Lipofectamine mediated RNA aptamer delivered and Wound scrap assay in cell lines confirms that the synthesized 18mer oligo has significant molecular level inhibition activity toward cyst formation and spreading in poly cystic condition in ovary.

## Introduction

Aptamers are primarily nucleotide sequences with high specificity towards target molecules. In recent times, they are successfully explored as better therapeutics to treat diseases and disorders. Time-consuming and low throughput procedures have been in practice to design and synthesize aptamers *invitro* (1). Therefore, in silico non-SELEX approach is the sagest choice to perform the selection of aptamers, which involves the construction of oligonucleotide library without amplification and binding them with suitable target protein unlike SELEX (2). Designing the RNA aptamer for the validated biomarker will help us to normalize the disease state at genetic level. Hence, the abnormal translation of target gene can be controlled by delivering a well-designed aptamer for the Response Elements (RE) of drug target. Response Elements are the key elements involved in the activation of target gene regulation. We have pinpointed RE specific for the regulation of abnormal expression of drug target. Further, iterative docking of aptamer with drug target helped us to identify the necessary base pairs for inhibition studies.

Biomarkers are medical signs for the persistence of certain illness or disease so inhibiting these biomarkers for specific pathophysiological condition at molecular level will be a better drug target choice to superintend the disease (3). Any alterations, such as genetic mutations, altered protein status, altered metabolites supports in inferring the potent drug target of disease. Validation of drug target is one of the requisite procedures in drug discovery and the criteria for validation depend on the certainty of the clinical end point. Since the exact cause of Poly Cystic Ovary Syndrome (PCOS) is imprecise (4), it is tedious to identify the best drug target among them clinically. Subsequently, system biology being an emerging field can be the effective way to detect and analyze the molecular targets of PCOS. Previously, researchers have found out that 500 biomarkers are prevalent in PCOS (5). In this research work, we used Protein Protein Interaction Network (PPIN) to select a highly influenced biomarker to be a drug target and also, we have efficiently employed other system biology tools to prioritize them. Finally, we have constructed the aptamer library for drug target through Non-SELEX fragment approach correspondingly assessed its affinity and stability.

## Materials and Methods

### Functional Network of PCOS Biomarkers

Polycystic ovary syndrome biomarker details and analysis of their interactions were done using DisGeNET and Cytoscape respectively. DisGeNET platform comprises a wide range of scientific information on Gene Disease Associations (GDA) and also facts from scientific literature by various text excavating methods (10). We have reclaimed the list of genes expressed by the query of umls: C0032460. Proteins of all the obtained genes were used to construct a functional Network with the help of STRING (11). The level of function confidential has set as 0.7 (medium) to prioritize interactive nodes (Figure 1A).

**Figure 1.**
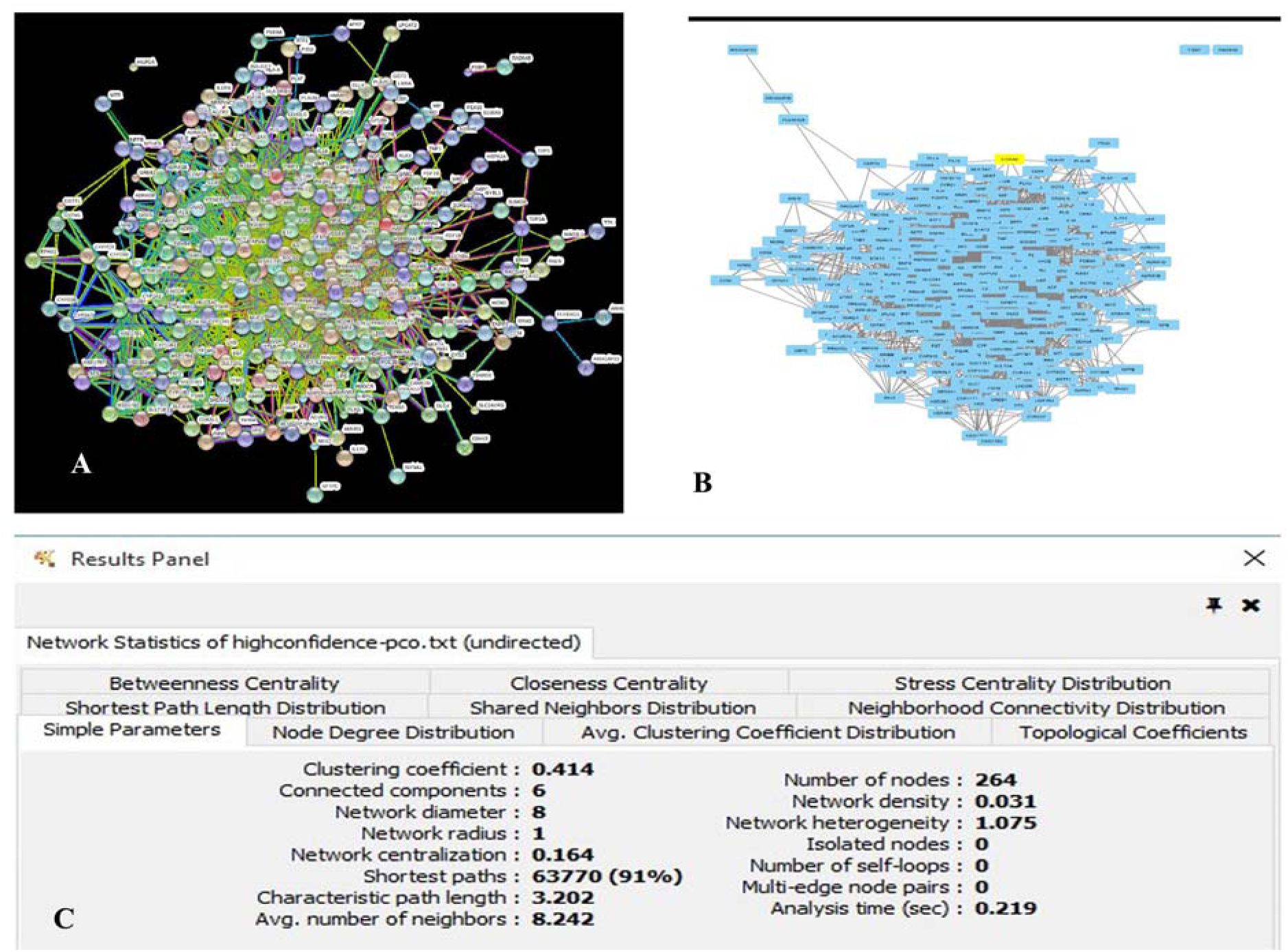
Network analysis of functional proteins, (1A) Protein-Protein Interaction Network of PCOS, (1B) Cytoscape view of constructed network, (1C) Undirected Phylogenetic layout properties

### Screening of target Nodes (Proteins)

Centrality measurement is a novel statistical tool to analyze the crucial nodes in a huge network. Cy1to NCA of Cytoscape 3.2 was employed efficiently to visualize and analyze the filtered network (12). The net statistical properties of constructed network were analyzed (Figure 1B) (13). To define the specific target from a dense network, the nodes interfering the pathways, number of neighboring nodes and the distance between them were considered as important criteria. We adopted three types of centrality analysis among eight, to sort the highly interactive nodes in the constructed network such as Betweenness, Closeness and Shortest path length. Significance of protein interactions without consideration of their weightage yielded desired result. At last, we have utilized two relative centrality analyses (ie, closeness and betweenness) that sorted 10 proteins, which were topologically and functionally relative respectively (Table 1). Self-loops of nodes were removed to avoid duplicate scores.

**Table 1.**
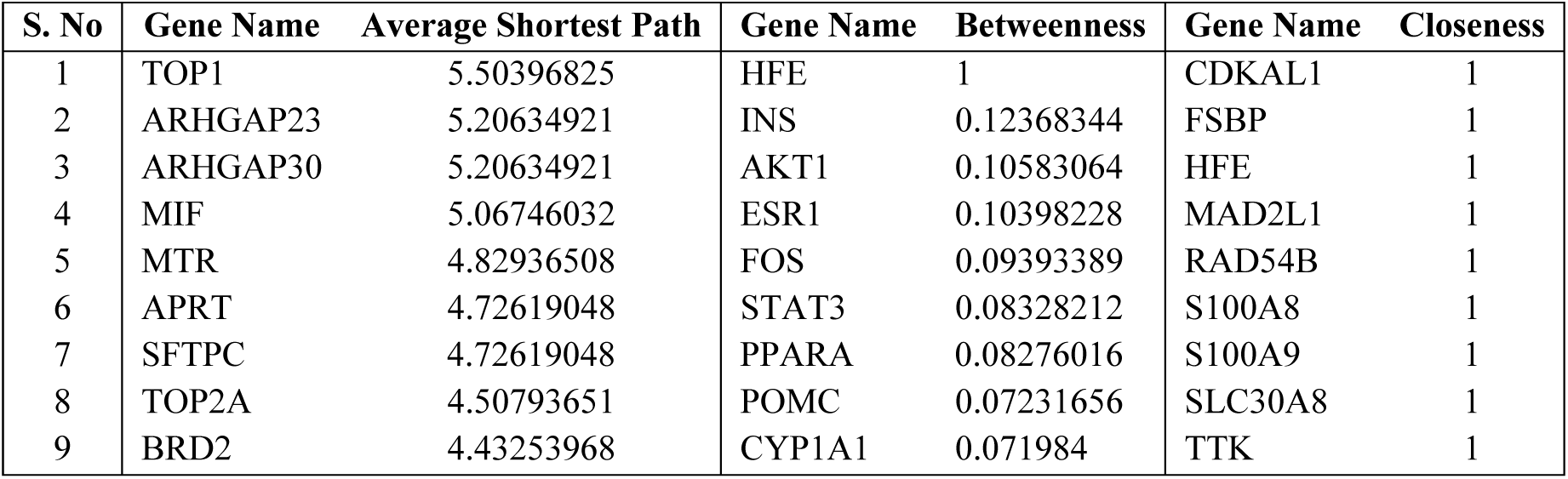
Selective scored Genes.

### Druggability analysis on identified Candidate

Druggability analysis has been used to predict the receptiveness and stability properties of a biomolecule regarding xenobiotics, which were curated to prioritize drug targets. The number of pockets, druggable score and pocket volume present in the target protein can be identified by physicochemical and geometric properties necessary to determine the efficiency of the target for designing aptamer accordingly. Binding site prediction, their analysis and druggability assessment of target protein based on protein heavy atom coordinates using Support Vector Machines (SVM) were predicted by DoGSite Scorer of University of Hamburg (14). Assessment of pocket volume, lipophilic character and pocket enclosures were used for simple score calculation and suggests the capability of a protein to act as a drug besides their functional mechanism (14). We have used the structure of target protein (PDB ID: 5HLV) which contains zinc and calcium bound complex. Structural analysis shows that eight chains containing 804 residues were interpreted. Based on the functions of refined proteins, our drug target was chosen (Table 3). On analyzing the functions of selected proteins from BioGRID 3.4, the pathway found is promising from others which may affect the normal functioning of human system.

### Selection criteria of Regulating Elements

Functional elements such as Response elements (RE), Exons, and Transcriptional Factors were investigated for their ability to influence the suppression of cyst formation in ovaries. By extensive literature survey, we have uncovered the general mechanism behind the activation or regulation of proteins in both molecular and cellular level (Figure 4). S100A8 Response elements binding sequences located in the locus 1q21 of chromosome were distinguished in the promoter region of S100A8 to construct mRNA analog library, employed to find all possible expression governing elements in upstream using Transcriptional Regulatory Element Database (TRED) (15). In addition to this, it is fundamental to know the entire detail of transcriptional factors that exclusively bind to S100A8. PROMO, an online tool which unraveled Transcription Factor Binding Sites (TFBS) in DNA sequences of species of interest (16). It has been applied to find all the TFBS in the upstream promoter region (upto1Kb) of S100A8. The result was rationalized by network analysis of S100A8. RE are the inducers of the receptor and ligand interaction which results in the expression or activation of a particular protein. Since this is the key element for the control of expression of target, it is chosen for the modelling of RNA against target protein. Glucocorticoid Response Element (GRE) (8), Hypoxia Response Element (HRE) (17), Antioxidant response element (ARE) (18), Interferon Gamma (INF-γ) Response Element (IRE) (19) influences the target gene as a response element.

### Glucocorticoid Response elements (GRE) for S100A8

Glucocorticoid Response elements, class of nuclear receptor family involves in signaling of small molecules that have the proficiency to influence the downregulation of our target protein through regulation of leukocyte transmigration (20). Glucocorticoids induce genes like macrophage migration inhibition factor ultimately resulting in downregulation of inflammation of cyst (20). Earlier research report shows that GRE consists of two half sites with three base pairs spacer and its consensus pseudo palindromic sequence is 5’ C<u>AGAACA</u>**TCA**TGTTCTGA 3’ (20).

### Construction of fragment library

RNA Composer, utilizes the Dot-Bracket format notation of the given secondary structure sequence where it models the RNA based on the canonical overlapping base pairs and 3D element is chosen from RNA frabase database (21). The consensus sequence was segregated as fragments in such a way that six nucleotides at a stretch was considered for RNA analog library construction. This was later utilized for binding with target by RNA-Lim method to recognize the various conformations exhibited in protein (22). This is modelled to achieve highly précised sequences. Diversity in the exhibited conformations of ssRNA-protein complexes were meticulously sampled by constructing fragment library. MC-Fold | MC-Sym pipeline was employed to obtain secondary and tertiary structure of constructed RNA Aptamer (23).

### Affinity and Stability studies

Refinement of docking results for their chain forming poses minimized the difficulty of designing an aptamer (24). Fragment based approach was adopted for competent docking with protein (25). This methodology is unusual and has numerous advantages over conventional rigid based docking (25). PatchDock, an freeware for effective protein protein docking using transformation (26). Based on the global binding energy, FireDock refines the docking result as top 10 proteins. It deciphers the result by flexible refinements rather than rigidity of protein and also optimizes side chain residues, minimizes the rigid body conformation (27). Previously, it was reported that the stability of RNA is checked by identifying inverted repeats that form stable hairpin loops for the particular nucleotide sequence (28). Since Oligoanalyzer, an inclusive oligonucleotide scrutinizer checks various properties for the given input nucleotide sequence, it was analyzed for hairpin loop (29).

### Wound Healing assay

MCF-17 cell lines were seeded into plate and incubated for 24 hours. Sterile microtip was used to make scratch on plate. 20 µl of Sample in Lipofectamine^TM^ 2000 was dissolved in DMSO and loaded into the plate. The results were seen after 4 hours of treatment. Cell migration Inhibition effect of aptamer was compared with control plate.

## Results

### Curation of functional proteins

As initiation in designing an aptamer, we have recovered 434 genes participating in PCOS, from which STRING annotated 267 branching nodes are as first shell of interactors. The obtained network consists of 1788 edges, 267 nodes with average node degree of about 13.44, also has clustering coefficient 0.48, which ultimately indicates that the network is significant. Clustered nodes in the network indicate the proteins connected with greater interaction. However, these nodes have functions in common pathways. Normally, relationships between nodes are considered as strongly orthologous (Figure 1A). The implication of S100A8 as an important node in the network epitomizes proteins with known or predicted structure respectively. It is notable that the presence of colored nodes of our network specifies co-occurrence, gene fusions in disease condition where highly interactive nodes were segregated from rest (Figure 1B) that benefits in elimination of genes with poor interaction. FSBP, RAD54B, HILPDA are the only proteins unconnected with the network.

### Assortment of Highly scored druggable proteins

Though, the complex network is tough to investigate, centrality analysis ranks the network elements based on their significance in interaction. We have employed undirected network to build the disease associated network which inferred us the functional associations among the nodes (Figure 1C). Towards this, we explored the statistical properties to categorize efficiently the criteria for druggability (Table 1). Properties such as Betweenness centrality, Closeness centrality, Stress centrality, Shortest path length distribution, shared neighbors distribution, neighbor connectivity distribution, simple parameters, node degree distribution, average clustering coefficient distribution and topological coefficients were analyzed in Cytoscape. It was evident that some of the genes have greater and stronger interaction with one another which have feasible probability of targeting PCOS for its treatment. One of the statistical properties of the network exerts clustering coefficient of 0.414, which indicates the clustering of nodes and number of self-loops were zilch (Figure.1B). As depicted from Table.1, we understood the statistical property of the genes whose average shortest path length ranges between 5.50396825 - 4.43253968. Genes enlisted in Betweenness ranges between 1 – 0.071984. However, entire genes related to PCOS were selectively scrutinized as top 10 from the rest. As mentioned in Table 1, significance of closeness and Betweenness was helpful in topological analysis of the scattered network. The path length graph shows the average length of 3 have maximum frequency of ≥26,000. In Betweenness centrality 20 neighbors intensively accumulate at 0.02 - 0.05. In closeness centrality, the nodes ranges from 0.35-0.45 and in twentieth neighborhood the clustering of nodes is high compared to another neighborhood.

### Target Compatibility Evaluation

Drug pocket properties, structure availability in PDB and functional relevance in PCOS are the criteria we have adopted to prioritize the target candidate, in which S100A8 shows better average evidences than other proteins (Table 2 and 3). Details of potential binding pocket details based on the 3D structure (1XK4) of S100A8 is given in Table 4. Also, by finding number of drug pockets we have chosen S100A8 as drug target. Considerably, we have inspected Response elements specific for S100A8 in order to down regulate the cell migration phenomena in cystic ovaries (6). This calcium binding protein (S100A8) acts as a ligand for Receptor of Advanced Glycation End Products (RAGE). S100A8 causes uteroplacental perfusion deficiency which leads to embryo abortion that supports the competence of our target selection (7). The expression of S100A8 is regulated by many REs but GRE is effective among them (8). S100A8, a member of S100 family that generally involves in signal transduction. Structural analysis shows our drug target has two helix loop helix Ca2+ binding domains known as EF hands and exists as complex with S100A9. The concentration of calcium affects the signaling capacity of S100A8. Calprotectin is present in 1q21 locus of chromosome 1 in human and has a molecular weight of 10-12 KDa. During tumor development, chromosomal rearrangements takes place in the locus of S100A8 gene, this majorly contributes to the cyst formation in PCOS. Also, serum Calgranulin (S100A8 and S100A9) levels are higher in women with PCOS than normal women (9). This evidently shows that binding to RAGE facilitates S100A8 to perform p38 MAP Kinase signaling through calcium phosphorylation, leading to cell migration eventually causing cyst formation.

**Table 2.** Structural and Druggable screening of selective proteins.

| Gene | Protein Name | PDB ID | No. Pockets | High Scored pocket | Drug Score |
| --- | --- | --- | --- | --- | --- |
| S100A8 | S100 Calcium Binding Protein A8 | 1XK4 | 21 | P_0 | 0.8 |
| MAD2L1 | MAD2 mitotic arrest deficient like | 1GO4 | 21 | P_2 | 0.86 |
| PPARA | Peroxisome Proliferator activated receptor alpha | 2REW | 5 | P_0 | 0.81 |
| STAT3 | Signal Transducer & Activator of transcription 3 | 4E68 | 21 | P_1 | 0.75 |
| POMC | Proopiomelanocortin | 4Z2A | 11 | P_2 | 0.66 |
| PIK3R1 | phosphoinositide 3-kinase regulatory subunit 1 | 4ZOP | 21 | P_0 | 0.8 |
| TOP1 | Topoisomerase 1 | 1A31 | 16 | P_1 | 0.82 |
| MIF | Macrophage Migration Inhibitory Factor | 3U18 | 10 | P_0 | 0.81 |
| MTR | 5 Methyltetrahydrofolate-Homocysteine S-Methyl transferase | 1Q8J | 21 | P_1 | 0.78 |
| ARHGAP23 | Rho GTPase Activating Protein 23 | 3PP2 | 8 | P_0 | 0.78 |
| ARHGAP30 | Rho GTPase Activating Protein 30 | 3BYI | 21 | P_1 | 0.81 |
| APRT | Adenine Phosphoribosyl transferase | 6FCH | 6 | P_2 | 0.86 |
| SFTPC | Pulmonary Surfactant-Associated Protein C | 5FFR | 4 | P_0 | 0.85 |
| TOP2A | DNA Topoisomerase II Alpha | 4FM9 | 21 | P_1 | 0.81 |
| BRD2 | Bromodomain Containing 2 | 5UEW | 5 | P_0 | 0.81 |
| HFE | Homeostatic Iron Regulator | 1A6Z | 21 | P_0 | 0.82 |
| INS | Insulin | 1FGY | 8 | P_0 | 0.64 |
| AKT1 | AKT Serine/Threonine Kinase 1 | 4EKL | 16 | P_0 | 0.83 |
| ESR1 | Estrogen Receptor 1 | 2Q70 | 13 | P_1 | 0.84 |
| FOS | Fos Proto-Oncogene, AP-1 Transcription Factor Subunit | 1A02 | 16 | P_0 | 0.81 |
| CYP1A1 | Cytochrome P450 Family 1 Subfamily A Member 1 | 4I8V | 21 | P_0 | 0.82 |
| CDKAL1 | CDK5 Regulatory Subunit Associated Protein 1 Like 1 | 4Y72 | 21 | P_1 | 0.86 |
| FSBP | Fibrinogen Silencer Binding Protein | 1D7H | 7 | P_0 | 0.55 |
| RAD54B | RAD54 Homolog B | 1Z3I | 21 | P_0 | 0.8 |
| S100A9 | S100 Calcium Binding Protein A9 | 6DS2 | 21 | P_0 | 0.81 |
| SLC30A8 | Solute Carrier Family 30 Member 8 | 6C08 | 21 | P_0 | 0.81 |
| TTK | TTK Protein Kinase | 3DBQ | 11 | P_0 | 0.82 |

**Table 3.**
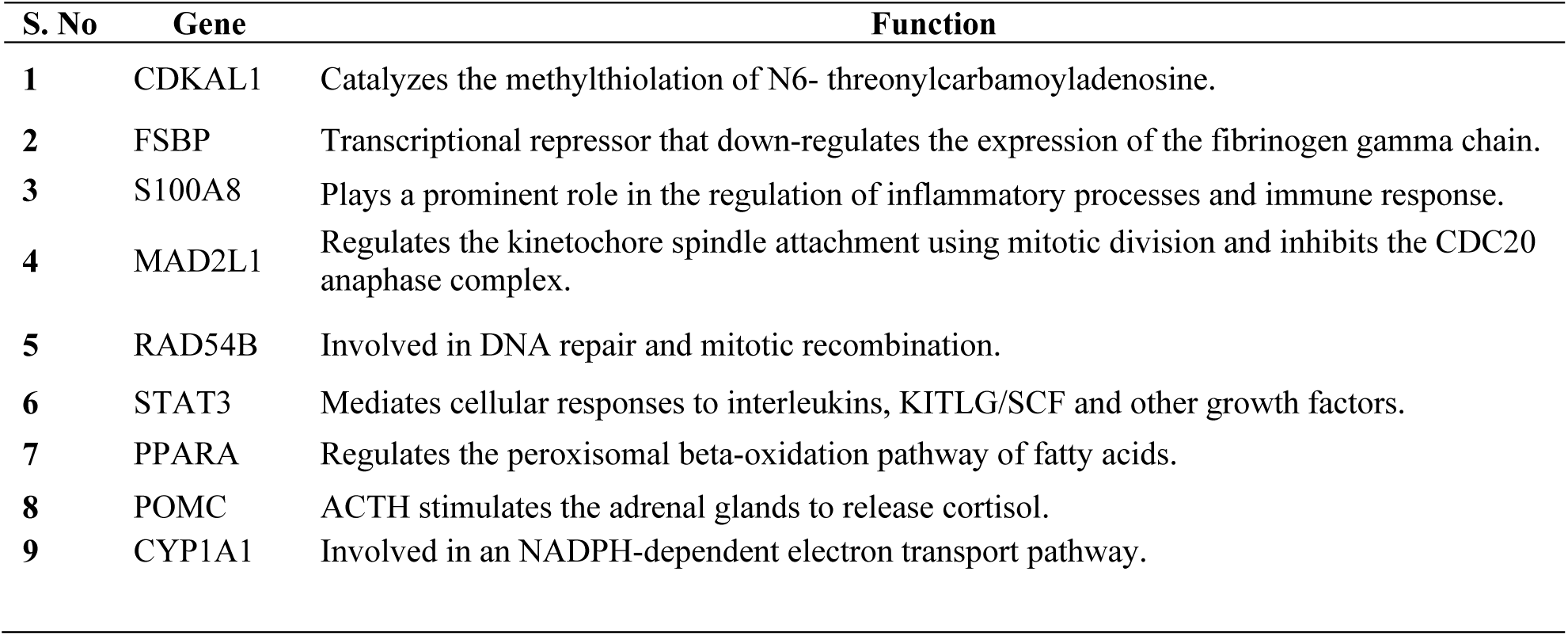
General functions of Prioritized Nodes.

**Table 4.** Druggabilty Assessment of S100A8.

| Name | Volume A<sup>2</sup> | Surface A<sup>2</sup> | Drug Score | Simple Score |
| --- | --- | --- | --- | --- |
| P_0 | 2307.87 | 2377.93 | 0.81 | 0.59 |
| P_1 | 2179.99 | 2093.63 | 0.81 | 0.65 |
| P_2 | 795.42 | 1549.19 | 0.78 | 0.51 |
| P_3 | 457.44 | 707.62 | 0.64 | 0.34 |
| P_4 | 209.65 | 195.14 | 0.66 | 0.0 |
| P_5 | 180.41 | 168.05 | 0.6 | 0.0 |
| P_6 | 173.21 | 283.31 | 0.35 | 0.0 |
| P_7 | 166.91 | 273.52 | 0.37 | 0.0 |
| P_8 | 156.79 | 332.75 | 0.35 | 0.02 |
| P_9 | 137.56 | 211.73 | 0.33 | 0.0 |
| P_10 | 130.58 | 204.55 | 0.28 | 0.0 |
| P_11 | 127.43 | 196.55 | 0.27 | 0.0 |
| P_12 | 120.91 | 239.32 | 0.27 | 0.0 |
| P_13 | 117.99 | 208.39 | 0.25 | 0.0 |
| P_14 | 116.19 | 209.36 | 0.22 | 0.0 |
| P_15 | 109.89 | 180.88 | 0.26 | 0.0 |
| P_16 | 109.89 | 259.12 | 0.16 | 0.0 |
| P_17 | 106.85 | 216.05 | 0.15 | 0.0 |
| P_18 | 100.78 | 249.25 | 0.14 | 0.0 |

### Construction of RNA Analog Library of GRE

Characterizing RE for S100A8 was a real challenge which even met the foundation of influencing the syndrome. As noted earlier, RE being an interacting sequence with the binding receptor of protein plays a key role in regulating the expression. We have con**s**tructed an RNA analog library of the specific RE glucocorticoid to mimic the action of actual RE. Fragment based approach yielded us better interaction with S100A8 in comparison with rigid based docking, a conventional docking strategy. Six nucleotide Fragmentation approach of RNA Lim method developed 18 fragments with the consensus sequence of GRE (Figure 2). Among the fragments, three (Frag6, Frag9 and Frag10) interactions (Table 5) are found in active domain of target protein with lowest global binding energy and the optimal structure in physiological parameters is showed in figure.5.

**Table 5.** List of 18 RNA Analog obtained through fragment approach along with their binding energy used for docking with target protein S100A8 and aptamer designing.

| Fragments | Predicted $\Delta G$ (kcal/mol) | Fragments | Predicted $\Delta G$ (kcal/mol) |
| --- | --- | --- | --- |
| Frag 1 | -16.24 | Frag 10 | -30.51 |
| Frag 2 | -8.55 | Frag 11 | -14.34 |
| Frag 3 | -16.52 | Frag 12 | -15.66 |
| Frag 4 | -51.85 | Frag 13 | -15.17 |
| Frag 5 | -18.46 | Frag 14 | -11.21 |
| Frag 6 | -31.71 | Frag 15 | -18.70 |
| Frag 7 | -16.54 | Frag 16 | -10.68 |
| Frag 8 | -13.17 | Frag 17 | -21.76 |
| Frag 9 | -38.76 | Frag 18 | -23.71 |

**Figure 2.**
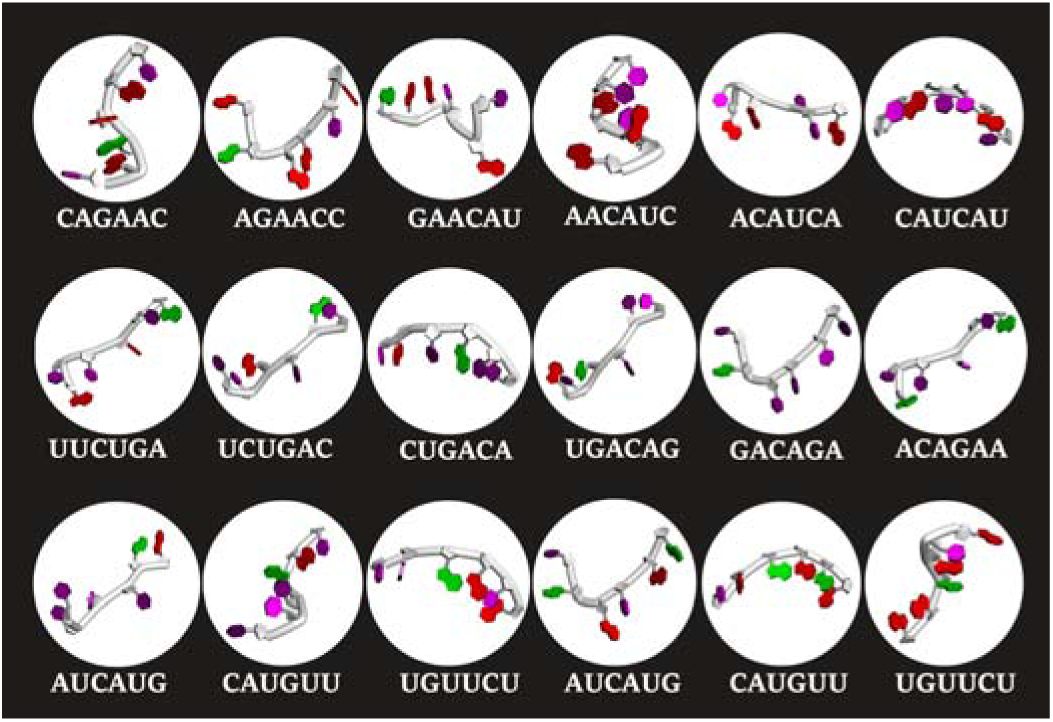
RNA fragment library by RNA-Lim method.

### Interaction and inhibition Analysis

Primarily, RAGE being the receptor for S100A8, was docked to confirm for its binding capacity in the appropriate domain, which effects in binding of the designed aptamers to inhibit the abnormality. As a result, we have found Arg 114 residue of A domain in RAGE interacted with Gln 44 residue in H domain of S100A8 by docking. To test the comparison of interactions with previous one, RAGE was later docked with GRE, whose binding energy is −24.38 comparatively grea**t**er than its binding with designed aptamers (−46.33) (Table 7). In Parallel, interactions of S100A8 with GRE and S100A8 with designed Aptamer were inspected for its binding capacity and found as −22.11 and −45.32 energy levels respectively. Since, the efficacy of aptamer is efficient at lower binding energy; the constructed aptamer is proven to be competent one (Figure 3).

**Figure 3.**
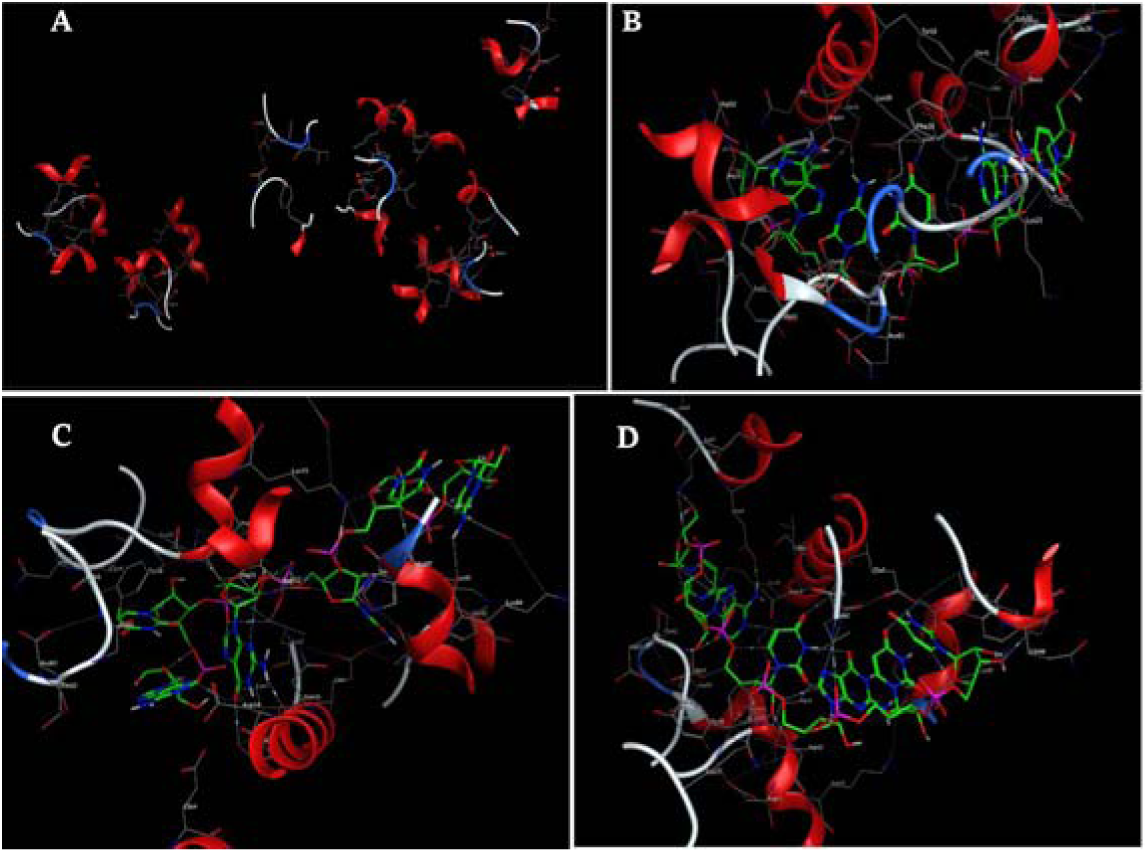
*Insilico* analysis. A) Active sites of S100A8, B) Binding pose of Fragment 6 with S100A8, C) Binding pose of Fragment 9 with S100A8, D) Binding pose of Fragment 10 with S100A8

**Figure 4.**
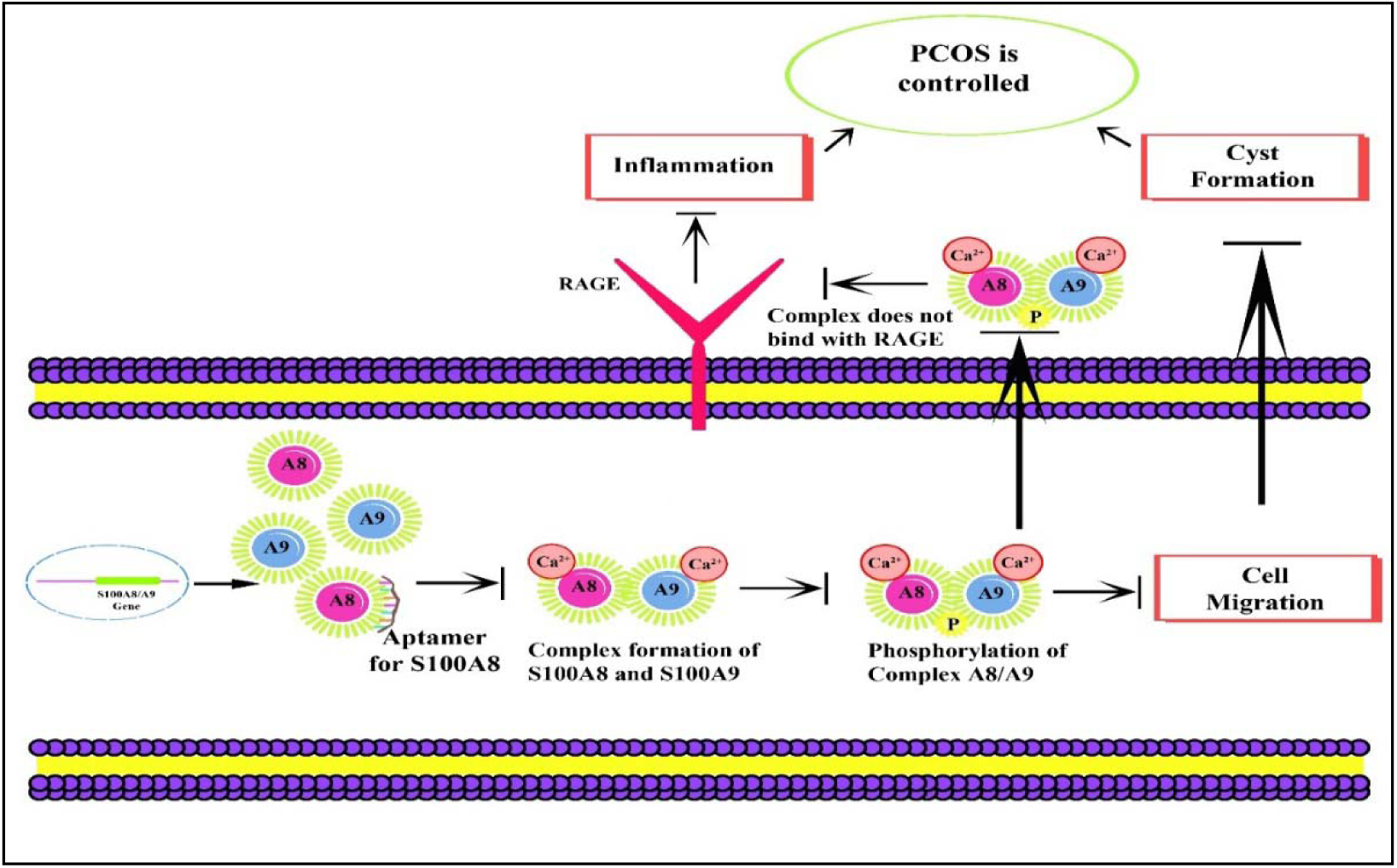
Mechanism of Response elements in cell migration process of S100A8.

**Figure 5.**
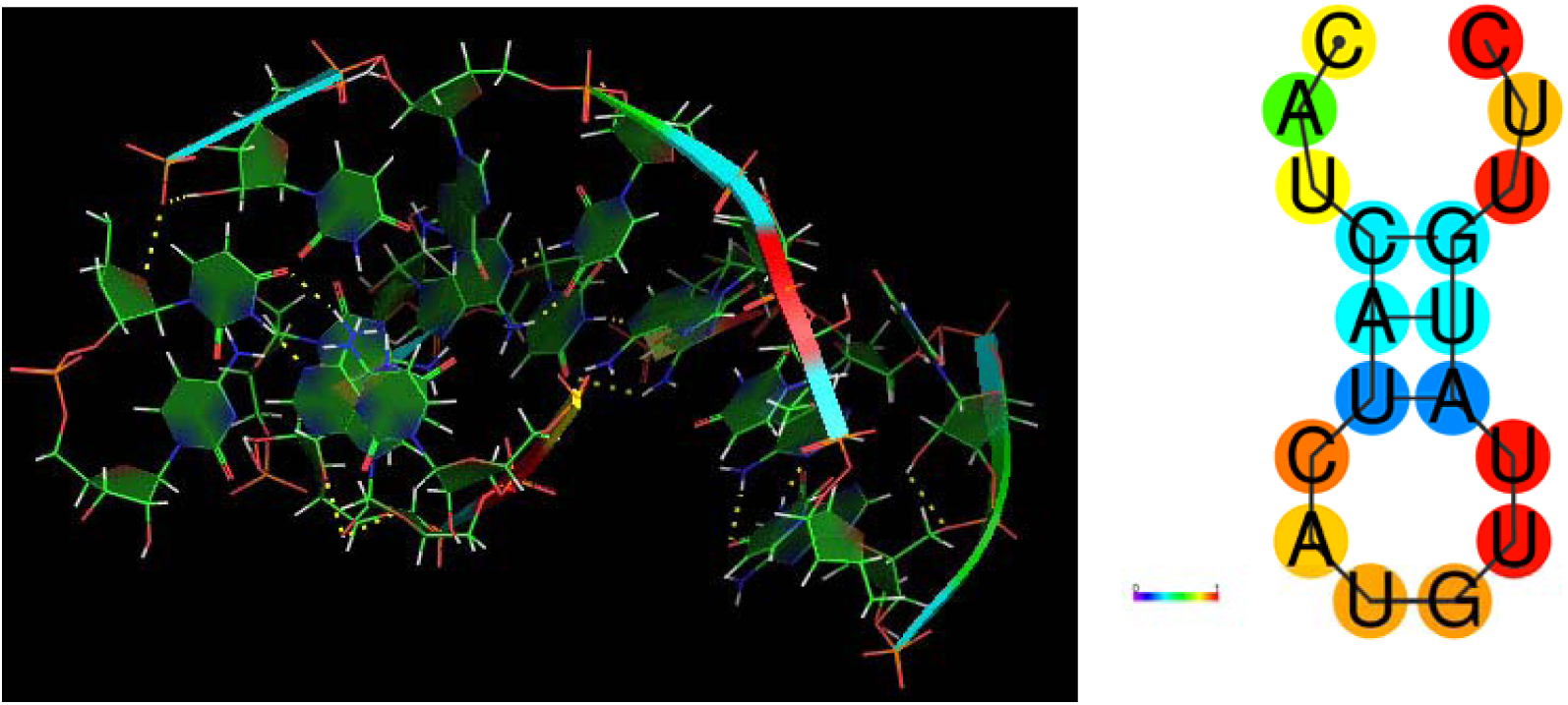
**a**. 3D Structure of designed RNA Aptamer with polar bonds. **B**. energy minimized structure at physiological pH.

### Stability of designed RNA Aptamers

Stability parameters comparison analysis among the Aptamers are stated in Table 6, in that Apt 1 has high stability at maximum of 41.8°C and also by system analysis, it requires minimum free energy (−27.93kcal/mol) for hairpin loop formation and possess low molecular weight (5327.4 g/mol) than other fragments.

**Table 6.**
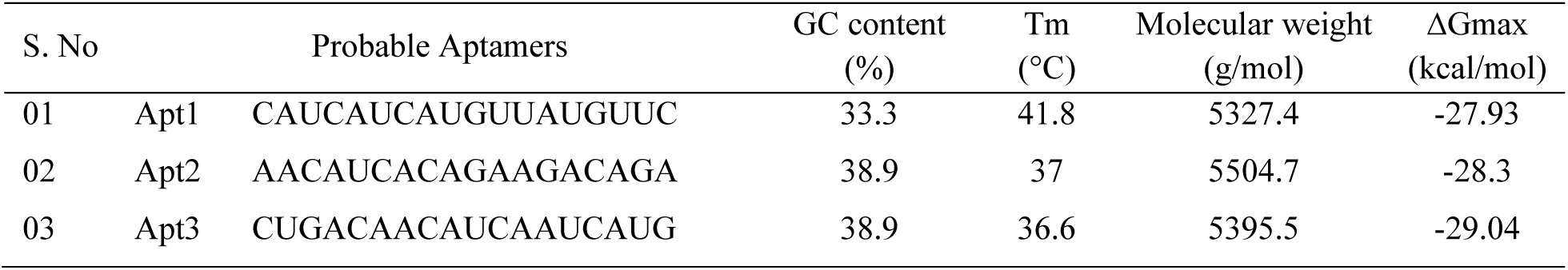
Enumeration of Aptamers whose stability and shelf life properties were listed above.

**Table 7.** Comparison of binding sites and energies of ligands with Targets

| S. No | Protein | Target | Binding Site | Global Energy |
| --- | --- | --- | --- | --- |
| 1. | RAGE | S100A8 | Arg A 114 → Gln H44 | -25.75 |
| 2. | RAGE | GRE | Arg B 203 → U<sub>14</sub><br>Arg B 228 → G<sub>17</sub> | -24.38 |
| 3. | S100A8 | GRE | Asn D 61 → C<sub>6</sub><br>Ala B 1 → U<sub>13</sub> | -22.11 |
| 4. | RAGE | Aptamer | Try B 118 → A<sub>13</sub><br>Arg B 216 → A<sub>13</sub><br>Arg B 218 → G<sub>10</sub><br>Asn B 25 → U<sub>16</sub><br>Gln B 24 → U<sub>16</sub> | -46.33 |
| 5. | S100A8 | Aptamer | Lys B 36 → U<sub>9</sub><br>Lys F 48 → U<sub>17</sub><br>Ser H 86 → A<sub>13</sub><br>Asp C 32 → G<sub>10</sub><br>Lys B 18 → G<sub>10</sub><br>Lys B 21 → G<sub>10</sub> | -45.32 |

### Anti-cell migration assay on MCF-17 cell line

In control plate, normal cells migrated and the 4mm wound scratched was covered within four hour of incubation but in the aptamer injected plate, there was no migration takes place when compared with control (figure.6).

**Figure 6.**
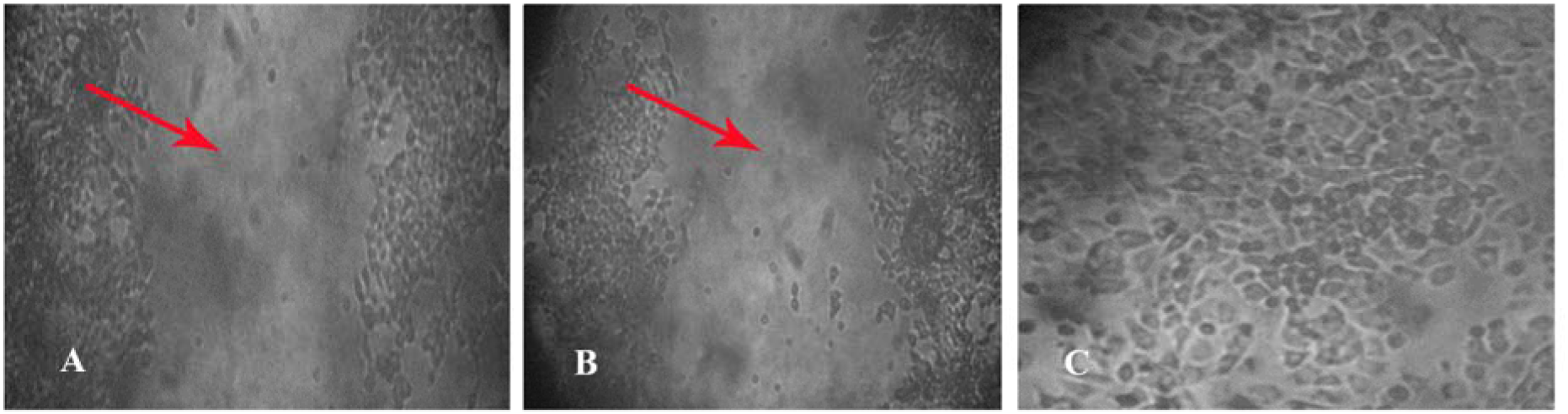
4mm scratch with sterile tip at zeroth hour (A), there was no cell migration observed (B) after fourth hour of incubation when compared with (C) control plate.

## Discussion

Though we removed three unconnected proteins in the disease network, the average connection of a single protein with other proteins is 6.696, and the clustering value also lead us to find the core node with respect to all possible druggable parameters. The high confidence interactome had only fewer strongly committed proteins, which would miss any better target. On the other hand, the low confidence network yields more unconnected nodes so the network construction we have chosen is of medium confidence level. Oligo fragments selected are by its binding ability on the active sites of discovered target but we obtained different binding site with least energy when they are compiled by18-mer fragments analysis. Priority of compiled fragment was given based on their binding energy but we haven’t checked the other possible orientation of fragments to construct optimal one. The effect of PCOS with and without the presence of Response elements. Binding of designed aptamer to S100A8- S100A9 complex leads to inhibition of cell migration process. Since the constructed aptamer showed better results than the Response elements, the positive effect on arresting cell migration shown in cell line study is only a preliminary one and we are studying the specific mechanism of inhibition, as per the results we could conclude that Apt1 has the potential to inhibit calcium binding protein (S100A8) ligand and the receptor protein called Receptor for Advanced Glycation End Products (RAGE) to inhibit the cancer cell movement and also to facilitate utero-placental perfusion.

## Acknowledgments

- We cordially thank **Tamil Nadu State Council for Science and Technology** for funding and got awarded as a Best project in the overall state level project contest for the year of 2017.
- We acknowledge Kamaraj college of Engineering and Technology for rewarding us the overall best project in “Technovision-2017”
- We acknowledge Chemical Computing Group for providing free license on MOE Software

